# Engineering synthetic phosphorylation signaling networks in human cells

**DOI:** 10.1101/2023.09.11.557100

**Authors:** Xiaoyu Yang, Jason W. Rocks, Kaiyi Jiang, Andrew J. Walters, Kshitij Rai, Jing Liu, Jason Nguyen, Scott D. Olson, Pankaj Mehta, James J. Collins, Nichole M. Daringer, Caleb J. Bashor

## Abstract

Protein phosphorylation signaling networks play a central role in how cells sense and respond to their environment. Here, we describe the engineering of artificial phosphorylation networks in which “push-pull” motifs—reversible enzymatic phosphorylation cycles consisting of opposing kinase and phosphatase activities—are assembled from modular protein domain parts and then wired together to create synthetic phosphorylation circuits in human cells. We demonstrate that the composability of our design scheme enables model-guided tuning of circuit function and the ability to make diverse network connections; synthetic phosphorylation circuits can be coupled to upstream cell surface receptors to enable fast-timescale sensing of extracellular ligands, while downstream connections can regulate gene expression. We leverage these capabilities to engineer cell-based cytokine controllers that dynamically sense and suppress activated T cells. Our work introduces a generalizable approach for designing and building phosphorylation signaling circuits that enable user-defined sense-and-respond function for diverse biosensing and therapeutic applications.

## MAIN TEXT

Cells universally use protein phosphorylation signaling networks to adapt to chemical and physical cues from their external environment. In metazoan cells, these networks consist of multi-layered pathways that rapidly and reversibly convert signals detected by cell surface receptors into diverse responses such as cell movement, secretion, metabolism, and gene expression (*1–4*). The ability to design artificial phospho-signaling circuits that exhibit native-like signaling behavior, yet can be programmed with custom-defined input/output connectivity, could be used to create powerful biotechnology (*5*), including human cell-based therapeutics that autonomously sense and respond to specific physiological signals or disease markers on a fast timescale (*6–8*).

Despite this potential, phospho-signaling circuit engineering has lagged behind that of genetic circuits (*9–11*), where advances in both microorganismal (*10*) and mammalian (*12*) settings have been enabled by design frameworks that leverage the intrinsic modularity of promoters and coding regions (*13, 14*), as well as carefully benchmarked sets of genetic parts (*15–17*)—features that have facilitated scaling of circuit complexity and fine tuning of circuit behavior using predictive quantitative models (*18–20*). Progress to date in engineering phospho-signaling circuits has included development of two-component phosphorylation pathways as programmable sense-and-respond modules in bacteria (*21–23*) and mammalian cells (*24–26*), and complex fast-timescale phosphorylation circuitry in yeast (*27–29*). In human cells, rewiring of native phospho-signaling networks has been used to create compact therapeutic programs (*30*) and sense-and-respond circuits that connect surface receptors to transcriptional outputs (*31, 32*). However, frameworks have yet to be described in human cells that facilitate the *de novo* design of multi-layered synthetic phosphorylation circuitry with programmable input-output connectivity and signal processing.

We sought to establish such a framework by emulating fundamental design features of native phospho-signaling networks, which are organized as sets of interlinked “push-pull” cycles comprising kinase and phosphatase activities that mutually act on a protein substrate (*33, 34*) (**Fig. 1A**). Changes in push-pull phosphorylation equilibrium occur rapidly in response to input signals and are quickly reversed upon input removal, enabling cells to adapt to environmental changes on timescales of seconds to minutes. These features motivated us to design tunable, interconnectable push-pull motifs as elementary units for constructing synthetic phospho-signaling circuits. To accomplish this, we took advantage of the intrinsic structural modularity of signaling proteins, which are typically composed of discrete domains that either carry out catalytic function (e.g., kinase, phosphatase) or specify interactions with other signaling components (*35–37*) (e.g., PDZ, SH3). The recombination of catalytic and interaction domains is thought to be responsible for rewiring signaling network connectivity during evolution (*37*), and has proved to be a versatile tool for engineering synthetic signaling network linkages (*38*) to create pathways with novel input-output relationships (*39, 40*) or to introduce new information processing function (*30, 41*). Here, we reasoned that orthogonal interaction domains could be used to mutually direct kinase and phosphatase domains to act on protein substrate targets, thereby establishing synthetic push-pull cycles that operate separately from native signaling networks.

**Figure 1.**
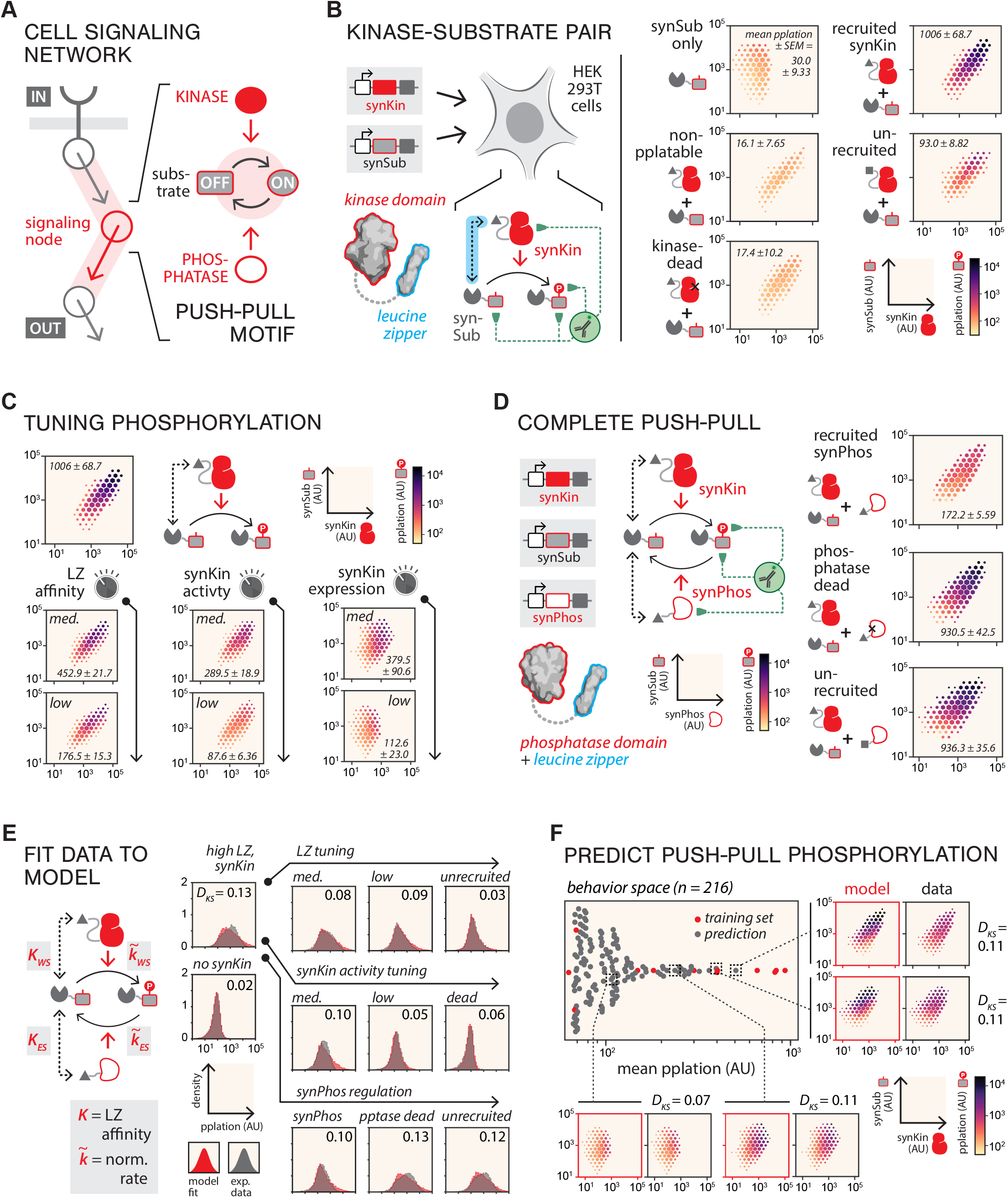
Building and tuning synthetic push-pull cycles in human cells. (**A**) Push-pull motifs, where kinase and phosphatase activities mutually regulate (red arrows) phosphorylation equilibrium (black arrows) of a substrate, are fundamental units that make up phospho-signaling networks. (**B**) Engineering synthetic kinase (synKin) and substrate (synSub) pairs. Plasmids (grey rectangles) encoding synKin and synSub proteins are transfected into HEK293T cells and measured for expression and phosphorylation via immunofluorescence flow cytometry (green dotted lines) after 36 h. Leucine zippers (LZs) mediate interactions (cyan dashed line) between synKin and synSub proteins (left). Expression and phosphorylation data for synKin/synSub allele combinations are shown on the right as hexagonal-hit-and-heat (HHH), through the whole study the expression space was uniformly binned into grids, the hexagon size indicates the cell counts in each bin, the largest hexagon size represents highest cell density, while the smallest hexagon size represents lowest density, with hexagon sizes in between representing log10-normalized cell counts proportionally (**fig. S5**). Values associated with HHH plots are mean phosphorylation (AU) ± standard error of the mean (SEM) (n=3). (**C**) Tuning synSub phosphorylation. Phosphorylation was measured for synKin/synSub compositions featuring parts that tune LZ binding affinity, synKin expression level, and synKin activity. The top panel shows behavior of the default synKin/synSub from Figure 1B for comparison. Numbers associated with HHH plots, mean phosphorylation (AU) ± SEM (n=3). (**D**) Complete synthetic push-pull. LZ-recruited synPhos dephosphorylates synSub (left). Values adjacent to HHH plots of synPhos variants indicate mean phosphorylation (AU) ± SEM (n=3) (right). (**E**) Modeling synthetic push-pull phosphorylation equilibrium. LZ interactions (*K_WS_*, synKin-synSub affinity [*W,* “writer”; *S,* “substrate”]; *K_ES_*, synPhos-synSub affinity [*E*, “eraser”; *S*, “substrate”) and component activity (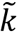_*WS*_, normalized phosphorylation rate; 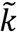_*ES*_, normalized dephosphorylation rate) of the push-pull (left), (see Model Description in the Supplementary Text). Push-pull data from Figure 1B-D were used for fitting. Model-predicted phosphorylation distributions (red) are plotted against experimentally measured distributions (grey). Kolmogorov-Smirnov divergence (*D*_KS_) values comparing model to experiment are shown for each plot. (norm., normalized) (**F**) Model-predicted push-pull behavior space. The beeswarm plot shows phosphorylation values for predicted (grey dots) and training (red dots) part compositions (216 total). HHH plots for compositions from across design space (indicated with black dotted lines) comparing predicted (red borders) and measured phosphorylation (black borders). *D_KS_* values comparing predicted and measured phosphorylation distributions are shown next to each pair of plots.

To validate this design strategy, we developed a protein domain part set that included engineered catalytic domains derived from components of immune phosphotyrosine (pY) signaling pathways (*42, 43*) (**figs. S1-2**). These proteins naturally utilize recruitment-dependent mechanisms of signaling activation (*44*) and are weakly expressed in most non-immune cell types (*43*). As an initial test, we fused various pY kinase domains (**fig. S3**) to leucine zippers (LZs)— small, highly specific heterodimerizing protein interaction domains with tunable interaction affinities (*41, 45*)—to create synthetically targeted kinases (synKins) (**fig. S4**). Synthetic substrate proteins (synSubs) for the synKins were constructed by fusing cognate LZs with ITAMs (immunoreceptor tyrosine-based activation motifs), conserved pY motifs involved in immune signaling pathway activation (*46*) (**Fig. 1B** and **fig. S4**). Plasmid constructs encoding pairs of epitope-tagged synKins and synSubs (**fig. S4)** were transfected into HEK293T cells, and multi-color flow cytometry was used to simultaneously measure component expression (staining against epitopes) and synSub phosphorylation (staining against phosphorylated ITAMs) in single cells (**fig. S5**). To optimize synKin function, we tested numerous kinase domain boundaries (**fig. S6**) and point mutations (**fig. S7**). We identified variants that showed strong expression and demonstrated phosphorylation activity toward synSub that was highly dependent on LZ-mediated kinase domain recruitment, as evidenced by non-binding, catalytically-dead, and unrecruited (non-cognate LZ) controls all showing little or no phosphorylation (**Fig. 1B**, right and **fig. S8**). Additionally, we demonstrated that expression of the synKin/synSub pair does not impair cell viability or limit cell growth (**fig. S9**).

We next tested whether we could tune the activity of synKin toward synSub by altering the molecular properties of our domain parts. We constructed several sets of synKin variants: LZ sequence variants were introduced to tune binding affinity to synSub (*45*), catalytic turnover rate was adjusted via previously reported pY kinase domain active site mutations (**fig. S4**) (*47*), and expression level was tuned by introducing Kozak sequence variants to synKin expression constructs (**fig. S1**), resulting in differential rates of protein translation (*48*). When we tested each synKin part set, we observed modulation of synSub phosphorylation across a 10-20 fold range (**Fig. 1C**). To create complete push-pull cycles with reversible synSub phosphorylation, we developed synthetically targeted pY phosphatases (synPhos). We took a similar approach as was used for the synKins, identifying domain variants derived from pY phosphatases involved in immune signaling (*49*) (**figs. S10-11**) and fusing them to the same LZ species as the synKin. As our data show, when co-expressed with a synKin/synSub pair, synPhos dephosphorylates synSub in a recruitment- and phosphatase activity-dependent manner (**Fig. 1D**). Taken together, these results validate our design strategy for constructing synthetic push-pull cycles and demonstrate that a simple part set consisting of catalytic and interactions domain variants can be used to rationally control intracellular phosphorylation equilibrium.

We next developed a model to quantitatively describe the relationship between push-pull equilibrium and part biophysical properties. To achieve this, we first converted single-cell fluorescence values into molecular equivalents for all push-pull compositions depicted in **Figure 1B-D** by normalizing different color fluorophores to an EGFP reference (**figs. S12-13**). These transformed data were then fit to a non-equilibrium thermodynamic model (**Fig. 1E** and **fig. S14**) to obtain part-specific parameters for LZ variant interaction affinities (*K_WS_, K_ES_*) and catalytic turnover rates of synKin and synPhos variants (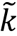_*WS*_, 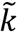_*ES*_) (**fig. S15**). The parameterized model was then used to predict push-pull phosphorylation levels for all part combinations within our design space (n=216 total compositions) (**Fig. 1F** and **fig. S16**). To validate these predictions, we constructed and measured compositions from across the predicted behavior distribution; all showed excellent overall agreement with the model (**Fig. 1F**), demonstrating that the functional modularity inherent in our design scheme lends itself to prediction of push-pull behavior based on individual parts properties.

Native signaling networks convert protein phosphorylation into molecular outputs through a variety of mechanisms (*50, 51*), including allosteric regulation of protein activity (*52*), changes in protein localization and stability (*53, 54*), and formation of new protein-protein interactions (*55*). In the latter case, phospho-specific binding domains recognize phosphorylated substrate motifs, forming interactions that facilitate downstream signaling (*56*). We hypothesized that circuit connections between our synthetic push-pulls could be engineered using SH2 domains, which bind to pY-containing motifs and are conserved among metazoans (*55*) (**Fig. 2A**). Prior to testing this, we used a transcriptional reporter (**fig. S17**) to not only validate synKin activity-dependent SH2/pY motif interactions, but also to identify part sets with orthogonal interaction specificities: tandem SH2 (tSH2) domains and an engineered multivalent SH2 respectively bound synKin-phosphorylated ITAMs (57) (**fig. S18**) a dual pY motif derived from Slp76 (**fig. S19**), with no observable crosstalk.

**Figure 2.**
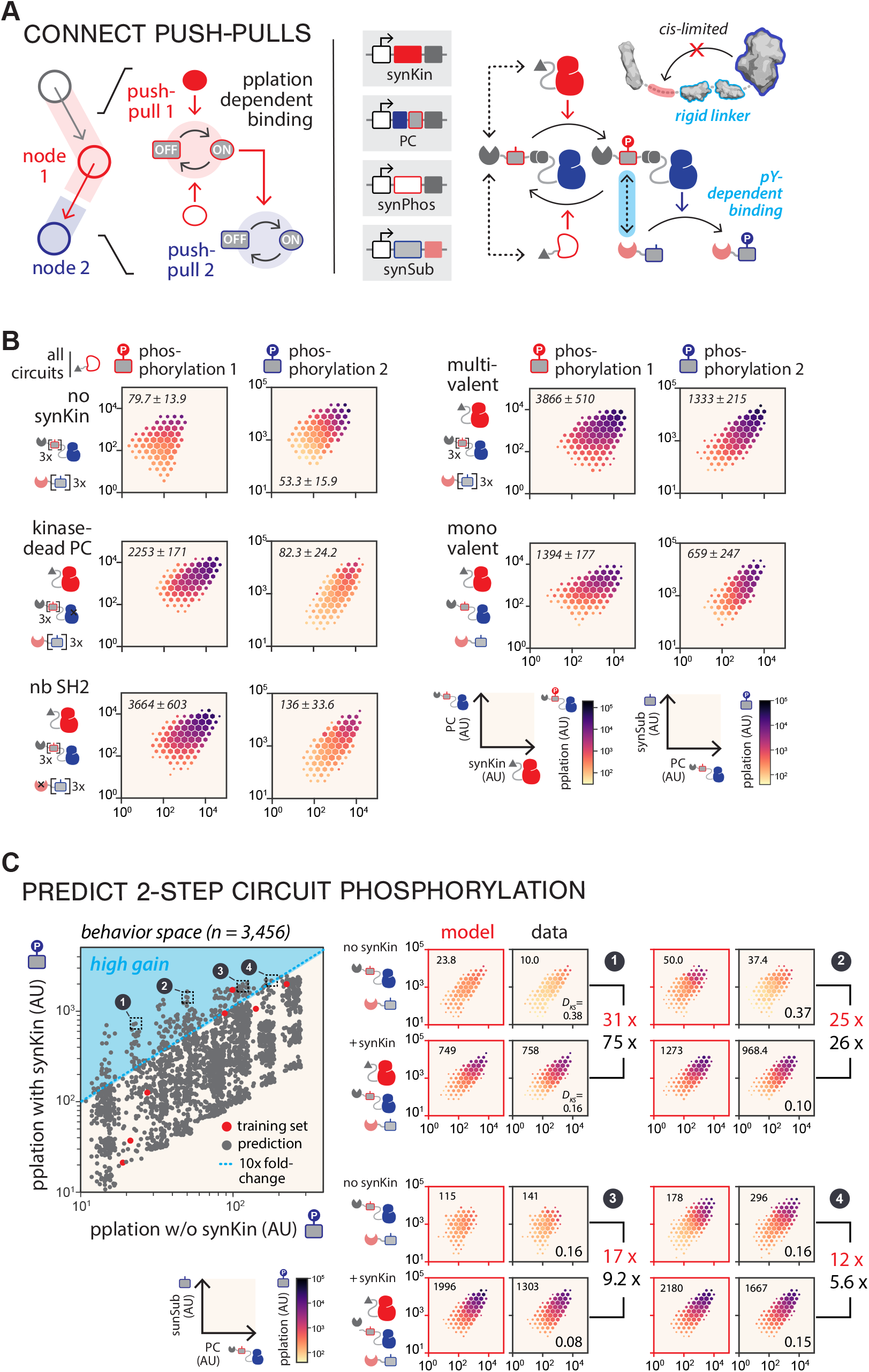
Building and tuning push-pull networks. (**A**) Two-step phosphorylation cascade circuit. To create a synthetic phosphorylation network connection, an upstream (red) push-pull cycle can be coupled to a downstream cycle (blue) using a phosphorylation-dependent interaction (left). The upstream synKin phosphorylates a “phospho-couple” (PC) protein, which functions as synSub for the upstream cycle and synKin for the downstream one (right). The PC contains a rigid linker domain (cyan outline) that prevents *cis* substrate motif phosphorylation while supporting SH2-mediated recruitment of a downstream substrate (cyan dashed line; upper right). (**B**) Two-step circuit validation. HHH plots for PC (left column) and synSub (right column) phosphorylation are shown for various circuit compositions. All circuit compositions contain synPhos. Brackets indicate the number of ITAM motif repeats. Values in each plot are mean phosphorylation (AU) ± SEM (n=3). (**C**) Model-predicted circuit behavior space for two-step circuit. The modeling framework was used to fit steady-state phosphorylation of two-step push-pulls from Figure 2B (**fig. S22**). Scatter plot shows predicted mean phosphorylation for compositions with (y-axis) and without (x-axis) upstream synKin (red dots, training set compositions; grey dots, model-predicted compositions [3,456 total]). Region of >10x fold-change (+/- synKin) is shown in cyan. Four high-gain circuits (indicated with black dotted lines) were constructed and tested for synSub phosphorylation. HHH plots for model-predicted (red border) and experimental measurements (black border) are shown. Values at the top of each plot indicate mean phosphorylation (AU) ± SEM (n=3) (right). *D_KS_* values are shown at the bottom of each experimental plot. Predicted (red) and experimentally measured (black) fold change values are shown to the right of the plots.

To test whether these phospho-dependent interactions could be used to link two push-pulls together, we engineered a “phospho-couple” (PC) protein that integrates the functions of an upstream synSub and downstream synKin by fusing a kinase domain to three substrate motifs and placing a rigid linker domain between them (**fig. S20**) to limit cis-phosphorylation (**Fig. 2A**, right and **fig. S21**). As our data show, this design facilitates sequential push-pull activation: when we expressed a 4-protein system (upstream synKin, PC, synPhos, and downstream SH2-synSub) in HEK293T cells, we found that phosphorylation of PC by the upstream synKin led to recruitment and phosphorylation of a downstream, tSH2-fused synSub (**Fig. 2B**). Sequential phosphorylation was dependent upon upstream synKin recruitment, PC activity, and SH2-mediated recruitment, as well as 3x substrate motif valency (**Fig. 2B**, right).

One important systems-level property of native phosphorylation cascades is their ability to stoichiometrically amplify weak input signals into macroscopic cellular outputs (*57*). To determine whether our two-step circuit architecture could be tuned to maximize amplification of an upstream input, we expanded our quantitative model to fit data from **Figure 2B**, obtaining part-specific parameters (**fig. S22A**) that allowed behavior predictions across two-step circuit combinatorial design space (n=3,456 compositions) (**Fig. 2C**, left). We identified a region of behavior space with compositions predicted to show a >10x fold-change in downstream synSub phosphorylation upon addition of an upstream synKin (n=261 compositions). Circuits from this “high-gain” region were enriched for features that are consistent with stoichiometric amplification, including a low PC:SH2-synSub ratio and strong synPhos activity (**fig. S23**). To validate model predictions, we selected several “amplifier circuit” compositions from this region to experimentally measure, demonstrating the general agreement of their behavior with model predictions (**Fig. 2C**, right). These results indicate that our part set and predictive modeling framework can be extended to guide the design of multi-push-pull networks with programmed signal-processing properties.

Having developed approaches for building, interconnecting, and predictively tuning synthetic push-pull motifs, we sought to engineer surface receptors that could couple extracellular ligand binding to changes in push-pull equilibrium (**Fig. 3A**, left). We constructed a pair of synthetic receptor scaffolds (**figs. S24** and **S25A**) consisting of flexible intracellular and extracellular linker sequences (**fig. S25B**) and transmembrane helices (**fig. S25C**). Kinase and LZ domains were appended to the cytoplasmic termini of the scaffolds, and FRB* and FKBP— domains that heterodimerize upon binding to the rapamycin analog AP21967 (*58*)—to their extracellular termini (**fig. S24**). We hypothesized that this architecture would enable ligand-induced synSub phosphorylation as a result of receptor dimerization-enforced proximity between the synKin and LZ (**Fig. 3A**, right). After transfecting a 4-protein system consisting of the receptor pair, synSub, and synPhos proteins, we observed a ∼20-fold change in phosphorylation upon ligand addition. Circuit induction was dependent on both LZ-mediated synSub recruitment and synKin activity. Additionally, we observed that elimination of synPhos activity resulted in a lower fold-change response (6.9x), demonstrating the importance of phosphorylation cycle reversibility for optimizing circuit performance (**Fig. 3B**).

**Figure 3.**
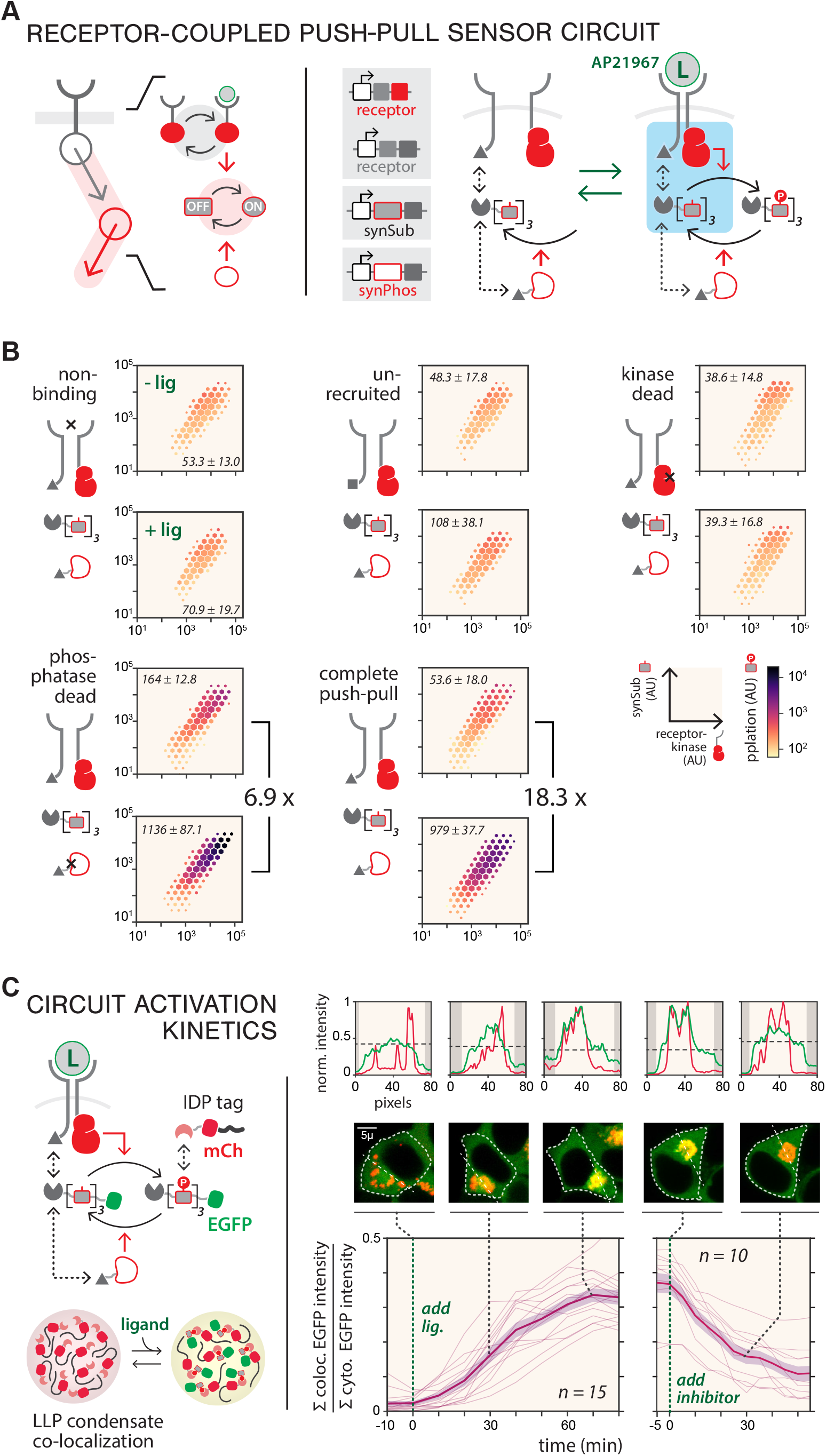
Constructing receptor-coupled push-pull sensor circuits. (**A**) Push-pull sensor circuit design. Reversible ligand binding to extracellular receptors triggers push-pull activation (left). The circuit is encoded as 4 genes on 3 plasmids; it includes two receptor chains (FRB*-TM-kinase domain and FKBP-TM-LZ), a synSub with 3 motif repeats, and a synPhos. Dimerization of extracellular FRB* and FKBP domains induced by ligand (AP21967) triggers colocalization of receptor-appended kinase with synSub (cyan box), leading to phosphorylation (right). (**B**) Testing receptor-induced phosphorylation. HHH plots depict flow-cytometry data from sensor circuit compositions analyzed 12 h after treatment with 200 nM ligand (+ lig) or a carrier-only control (- lig). Values in each plot indicate mean phosphorylation (AU) ± SEM (n=3). Phosphorylation fold-change values are next to each set of plots. (**C**) Measuring pathway activation using an LLP condensate co-localization reporter. The reporter consists of EGFP-tagged synSub and an mCherry-tagged SH2 domain tagged with an intrinsically disordered protein (IDP); phosphorylation leads synSub-EGFP recruitment to condensates and EGFP/mCherry colocalization (left). For activation experiments, cells were cultured for 24 h and then time-lapse images were taken every 10 mins following ligand addition over an 80 min time course to track pathway activation (left time course plot). For deactivation experiments 200 nM ligand was added for 90 mins followed by addition of the inhibitor imatinib mesylate (10 µM), and then time-lapse images were taken every 5 mins following inhibitor addition over a 60 min time course (right time course plot). Data are plotted as single cell trajectories (thin pink lines) for activation (n=15 cells) and deactivation (n=10 cells), with mean values (thick pink line) ± SEM (shaded pink bands). For selected time points (0, 30, and 70 min for activation and 0 and 30 min for deactivation), images of EGFP and mCherry (false-colored green and red, right middle) are shown for representative single cells, with the cell boundaries (dotted white outlines) as determined by custom segmentation software. Histograms (right top) show max-normalized EGFP and mCherry intensities along the straight white dashed lines drawn in images, and intensity is plotted for each channel in the same plot, with the black dashed line representing max-normalized cytoplasmic EGFP intensity; shaded regions, outside the cytoplasm. Scale bars, 5 μm.

To assess the timescale of activation for our receptor-mediated phospho-sensor circuit, we engineered a reporter system that allowed us to track the accumulation of phosphorylated synSub in real time using fluorescence microscopy (**Fig. 3C**). The reporter was created by fusing an SH2 domain and mCherry fluorescent protein to a PopZ tag (*59*), an intrinsically disordered protein that can sequester fused client proteins into cytoplasmically-localized liquid-liquid phase condensates (*60*) (**fig. S26**). This enabled us to monitor circuit activation by quantifying the co-localization of EGFP-fused synSub to the condensates as a proxy for phosphorylation (**fig. S27** and **movie S1**). We detected EGFP/mCherry co-localization within 10 mins following ligand addition and steady state was reached after ∼1h, while addition of a synKin inhibitor to the fully active pathway led to rapid synPhos-dependent de-localization (**Fig. 3C** and **movie S2**). For a circuit in which synPhos is absent, we found that de-localization occurs >10x slower, demonstrating that synPhos is critical for circuit reversibility (**fig. S28**). Fitting these data to a dynamic model (**fig. S29A**) yielded circuit activation and deactivation half-times of 28.8 and 22.8 min, respectively—rapid dynamics similar to those measured for cytokine signaling pathways such as RTK/JAK-STAT (*61*) and TGFβ/SMAD (*62*) (**fig. S29B**).

We next asked whether we could combine our phospho-sensor and two-step amplifier modules to create a sense-and-respond circuit capable of converting an extracellular input signal into expression of a transgene (**Fig. 4A**, left and **fig. S30A**). To promote membrane-to-nucleus signal propagation, we selected an amplifier composition (#2 from **Fig. 2C**) that showed both high gain and high downstream phosphorylation, and appended NLS and NES motifs to the PC to promote shuttling between the cytoplasm and nucleus. Regulation of transcriptional activation was implemented by fusing the second substrate to a synthetic zinc finger transcriptional factor (*20*) (synTF) to facilitate phospho-dependent recruitment of an SH2-fused transcriptional activation domain (TAD), resulting in initiation of EGFP reporter expression. We tested this 6-protein, 7-gene circuit in HEK293T cells and observed PC phosphorylation (13x fold-change) and EGFP expression (16x fold-change) in response to ligand addition (**Fig. 4A**, right). Non-NLS/NES-tagged (**fig. S30B**) and lower-gain circuit designs (**fig. S31**) showed <10x fold-change, underscoring the importance of shuttling and amplification as circuit design features.

**Figure 4.**
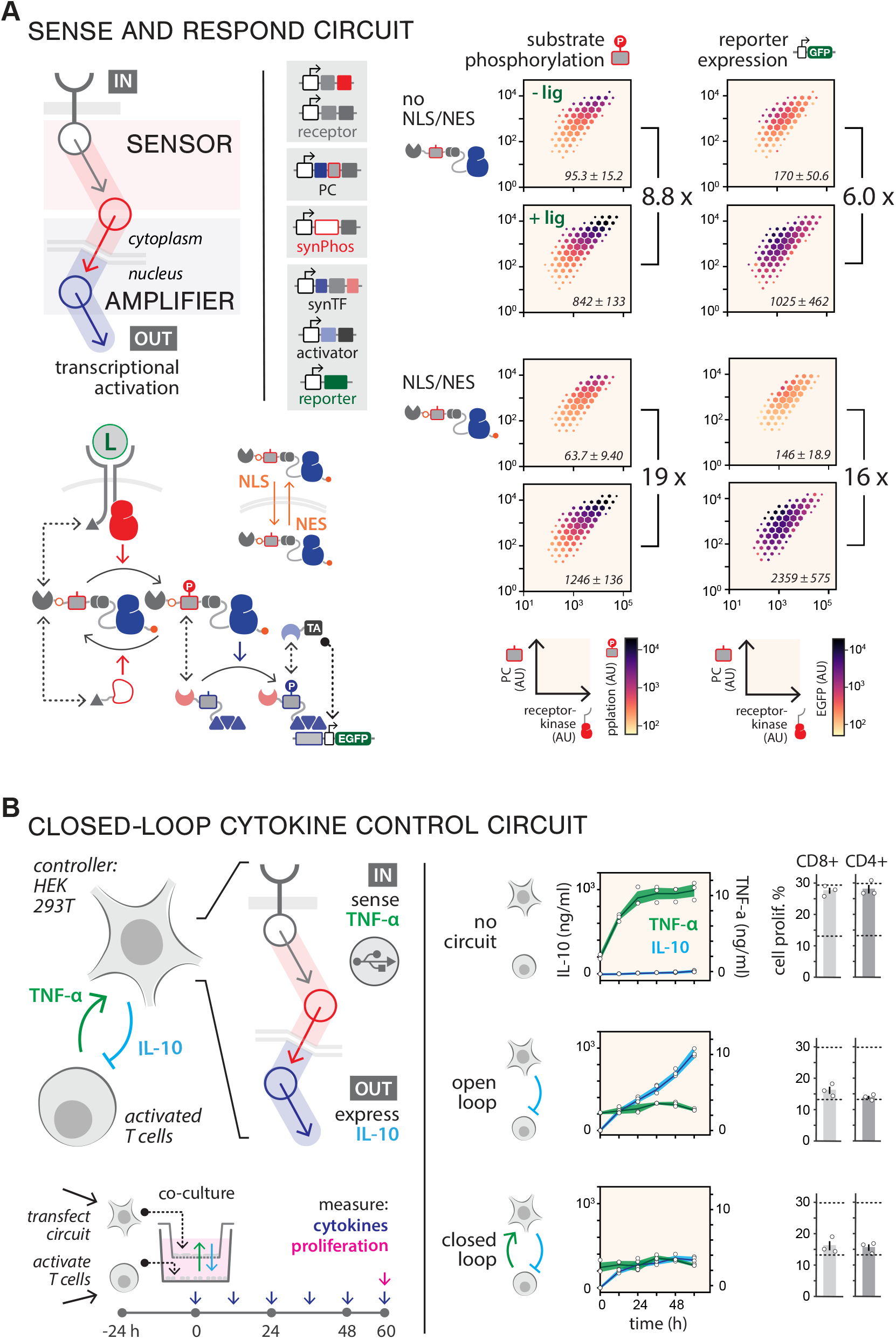
Phosphorylation circuit-mediated closed-loop sense-and-response function. (**A**) Using phospho-signaling to connect extracellular sensing to transcriptional output. The phospho-sensor circuit (Fig. 3) was coupled to a two-step amplifier circuit (Fig. 2), which was in turn coupled to a phosphorylation-dependent transcriptional reporter module (**fig. S19**), yielding a membrane-to-nucleus signaling pathway. The PC is tagged with nuclear localization (NLS) and nuclear export signals (NES) to enable shuttling between the nucleus and cytoplasm. HHH plots for PC phosphorylation (left column) and EGFP expression (right column) are shown for cells +/- ligand for compositions with and without NLS/NES tags. Values in the plots indicate mean fluorescence (AU) ± SEM (n=3). Fold-change values are shown to the right of each set of plots. (**B**) Engineering a phospho-signaling pathway for closed-loop therapeutic control. HEK293T cells expressing a circuit that can sense TNF-α (green arrow) and respond by secreting IL-10 (blue arrow) (top left) are placed in transwell coculture with activated T cells (bottom left) for 60 h, with media collected every 12 h to measure cytokine levels. T cell proliferation was assessed by EdU assay at 60 h. TNF-α and IL-10 time courses are shown for empty HEK293T cells (no circuit), constitutively IL-10 expression (open loop), and the sense-and-respond circuit (closed loop). Each circle represents a different PBMC donor (black line, mean values; shaded regions, ± SEM) (middle). Measurements of CD4+ and CD8+ proliferation, with circles representing data from 3 PBMC donors (error bars, indicating mean values ± SEM, n=3 measurements; upper dashed line, proliferation [maximum EdU signal] of activated T cells alone with no HEK293T cell; lower dashed line, proliferation of activated T cells inhibited with 500 ng/ml IL-10 [minimum EdU signal]).

As a demonstration of the translational potential of our framework, we engineered a circuit that senses TNF-α—a cytokine secreted by T cells that drives adverse inflammatory response (*63*)—and responds by secreting IL-10, a cytokine that inhibits T cell activation/expansion and TNF-α production (*64*) but has toxic side effects that limit its clinical utility (*65*) (**Fig. 4B**, left). We hypothesized that cells harboring this circuit could establish an anti-inflammatory control loop that suppresses T cell activation while maintaining both cytokines at low setpoints. We tested this by reconfiguring the sense-and-respond circuit in **Figure 4A**; we appended single-chain antibody fragments (scFvs) that recognize TNF-α (*66*) to the receptors (**fig. S32**) and replaced the EGFP reporter with IL-10. HEK293T cells equipped with this circuit were introduced into a transwell co-culture with CD3/CD28-activated human PBMCs, and cytokine production and T cell proliferation were assessed across a 60 h time course (**Fig. 4B**, left). For co-cultures containing cells with no circuit, we observed rapid accumulation of TNF-α and robust T cell proliferation (**fig. S33**), while cells constitutively expressing IL-10 (open-loop configuration) inhibited TNF-α secretion and T cell proliferation (**Fig. 4B**, right). Cells equipped with the sense-and-respond circuit (closed-loop configuration) also suppressed T cell proliferation but reached low steady-state levels of both TNF-α and IL-10 after ∼12 h. As indicated by modeling the dynamics of this system (**fig. S34**), this rapid setpoint convergence is likely dependent on the fast activation and deactivation rates of our phospho-signaling circuit, and may not be achievable with circuits that use molecular mechanisms that operate on slower timescales (**fig. S35**).

Here, we have engineered synthetic phospho-signaling circuits using a simple design logic in which push-pull cycles are utilized as building blocks, and circuit connectivity and information flow are defined through programmed protein-protein interactions. As we demonstrate, the composability of our design framework enables predictive tuning of circuit behavior and the use of nonequilibrium thermodynamic modeling to guide circuit design. While the part set we used in this study to demonstrate the practicability of our design scheme consisted largely of domains and motifs repurposed from native human immune signaling, our framework should facilitate incorporation of domains drawn from other sources or generated by computational design (*67*). Catalytic domains could be engineered to enhance circuit performance through activity tuning, or by introducing allosteric regulation (*68*). Since the functional specificities of our components are determined by recruitment, scaling to greater circuit complexity could be enabled by simply expanding the number of orthogonal interaction domains (*69*) in our part set. This could facilitate construction of circuit topologies that carry out advanced signal processing functions, such as Boolean logic enabled through multi-site phosphorylation (*70*), feedback connections that tune circuit dynamics or introduce ultrasensitivity (*71*), or multi-input-output circuits that can perceive and compute internal or external states (*72*).

Finally, because our circuits are post-translational and signal rapidly and reversibly, they can potentially support a broad array of cell-based diagnostic and therapeutic applications that require sensing of minute-scale physiological or pathological events (*8*). The plug-and-play configurability of our circuits should enable their coupling to diverse receptor inputs capable of sensing small molecules, bioactive factors, or disease markers (*73*). Because of their temporal responsiveness, the circuits may complement or offer advantages over other highly programmable circuit design schemes that signal by slower-turnover molecular mechanisms (e.g., transcription or proteolysis) (*74*). Additionally, since our circuits operate in parallel to native signaling pathways, they offer opportunities for programming signal-processing functions that are not possible for signaling circuits that harness native components to propagate signal. Furthermore, since they can be configured with human-derived protein domains and are relatively compact, circuits constructed using our design framework are likely to have low immunogenicity and could potentially be delivered to a diverse array of primary cell types to enable therapeutic sense-and-respond function.

## ACKNOWLEDGEMENTS

We thank members of the Bashor lab for helpful discussions. We acknowledge Ching-Hao Wang for his contributions to development of the modeling framework. We also thank Jared Toettcher and Steven Boeynaems for advice on engineering and monitoring protein condensates. We also thank Ryan Butcher for helping set up Nikon NASPARC imaging system for taking time-lapse movies. We also thank Siliang Li and John Her for helping build DNA constructs.

## Author contributions

C.J.B., N.D., and X.Y. conceived of the study. C.J.B. and N.D. carried out initial experiments. X.Y. built all DNA constructs and carried out experiments to generate the data published herein, with assistance from K.J., A.W, J. L., and J.N.. J.W.R., X.Y., and K.J. developed the quantitative modeling framework with input and supervision from P.M. and C.J.B.. X.Y. and A.J.W. developed the transwell assay supervision from S.D.O. and C.J.B.. K.R. developed image analysis and dynamic circuit modelling. X.Y., J.W.R., P.M., and C.J.B. analyzed the data. C.J.B., N.D., and J.J.C supervised the study. X.Y. and C.J.B. wrote the manuscript, with generous input from all authors.

